# Thermal plasticity of virulence differs between two latitudinally distant populations of the ash dieback pathogen

**DOI:** 10.64898/2025.12.22.695959

**Authors:** J.P. Soularue, C. Becans, A. Martelli, C. Lepoittevin, A. Aubert, Cécile Robin

## Abstract

*Hymenoscyphus fraxineus* is an invasive fungal pathogen responsible for the ash dieback epidemic, which continues to cause severe mortality of common ash (*Fraxinus excelsior* L.) across Europe. Following its likely introduction in northeastern Europe, the pathogen rapidly colonized most regions where common ash is present. As it spread southward, *H. fraxineus* encountered warmer climates and a higher occurrence of *Fraxinus ornus* L., a species resistant to the disease, which may have imposed novel selective pressures on *H. fraxineus* at the epidemic front.

Virulence is a key trait that is expected to evolve during epidemics and to exhibit plasticity in response to environmental variation. We investigated whether the plasticity of virulence in response to temperature and ash species has evolved in *H. fraxineus*. Using leaf inoculations, we characterized individual reaction norms for virulence in a long-established Lithuanian population and a recently established Italian population of *H. fraxineus*.

At moderate temperature, the two *H. fraxineus* populations showed similar levels of virulence on *F. excelsior*. Virulence decreased to a similar extent on *F. ornus* in both pathogen populations, indicating similar plasticity of their virulence in relation to ash species. Interestingly, increasing temperature reduced virulence in the Lithuanian population but did not significantly change the virulence expressed by the Italian population, which remained stable across temperatures. Although these findings warrant further investigation, they suggest that ash populations in southern Europe may be more vulnerable to ash dieback under warm conditions than previously anticipated.

## Introduction

The emergence of forest diseases caused by the accidental introduction of non-native pathogens has intensified markedly in recent decades (*e.g.*, Desprez-Loustau et al. 2010, Santini et al. 2013, Roy et al. 2014). These diseases inflict severe biodiversity losses on forest ecosystems and threaten the wide array of ecosystem services provided by natural and exploited forests (Keesing et al. 2010, Freer-Smith & Webber 2017, Guégan et al. 2020). Effective disease management strategies rely on accurate assessments of disease risk, which in turn require clear insights into the key drivers of forest epidemics. However, our understanding of these drivers remains limited, notably because forest disease management has historically overlooked the evolutionary dynamics shaping interactions between pathogens and their biotic and abiotic environments (Desprez-Loustau et al. 2016). Accounting for these evolutionary dynamics is becoming increasingly important as global changes facilitates the establishment and range expansion of invasive pathogens ( George et al. 2025)

Phenotypic plasticity refers to environmentally induced variations in phenotype ( Sommer 2020), typically represented using reaction norms (Ghalambor et al. 2007). In variable environments, plastic trait expression can give rise to polyphenism, a form of phenotypic diversity that may combine with genetic polymorphism (Mayr 1963, West-Eberhard 2003), and may sustain population adaptation to novel environmental conditions (Hoffmann & Sgrò 2011, Lande 2009, Merilä & Hendry 2014, Gomulkiewicz & Stinchcombe 2022, Vinton et al. 2022). The role of plasticity is especially critical in organisms such as some forest pathogens which have limited capacity to track favorable environments during the lifetime of an established genotype (Chevin et al. 2013, de la Mata & Zas 2023, Forsman 2015). Importantly, when phenotypic plasticity is heritable, the shape and diversity of reaction norms may evolve according to various patterns. Theoretical models predict that adaptive reaction norms can evolve when sufficient genetic variation in reaction norm shape (Via & Lande 1985) and substantial gene flow (Scheiner et al. 2012) are present. Such reaction norms confer high fitness to genotypes across environmental variation. For linear reaction norms, this evolution is expected to produce steep reaction norms when optimal phenotypes differ between environments (divergent selection), but flat reaction norms when stabilizing selection favors the same optimal phenotype across environments. This latter outcome, also called phenotypic buffering, is likely to arise for traits tightly linked to fitness, such as mycelial growth or the ability to colonize hosts in fungal pathogens, and is considered a special case of plasticity evolution (Bradshaw 1965, Eriksson et al. 2023, Reusch 2014). However, the evolution of reaction norms does not systematically lead to adaptive plasticity. Reaction norms may instead remain maladaptive and hamper invasion success when populations cannot track changing selective optima because environmental change is too rapid (Dupont et al. 2024), when environmental cues poorly predict future selective conditions (Ghalambor et al. 2007, Leonard & Lancaster 2020), when genetic variation underlying plasticity is limited (DeWitt et al. 1998), or when the costs of plasticity outweigh its benefits (Murren et al. 2015, Scheiner 2013). Understanding the invasive success of introduced forest pathogens, and anticipating their future epidemiological dynamics, thus requires understanding how plasticity evolve in fitness-related traits. Yet, empirical knowledge in this area remains limited (but see Garbelotto et al. 2015), making it critical to assess the diversity of individual reaction norms for fundamental traits in forest tree pathogens across plausible environmental variation.

Virulence is defined as the extent of damage a pathogen causes to its host (D’Arcy et al. 2001, Sacristán & García-Arenal 2008). In plant pathology, this quantitative component of pathogenicity is commonly referred to as aggressiveness (Pariaud et al. 2009), and typically quantified using measures as lesion size (*e.g.* Lygis et al. 2017, Kosawang et al. 2020) or symptom severity (*e.g*., Orton et al. 2019). Virulence is expected to evolve during the course of epidemics along complex and sometimes antagonistic trajectories (Sackett and Mundt 2005, Cressler et al. 2016, Dutt et al. 2022). However, excessively high virulence may reduce transmission by shortening the infectious period or limiting host availability. Under the virulence-transmission trade-off hypothesis (Anderson and May 1982), virulence is expected to evolve towards an intermediate optimum that balances within-host competition, infected host survival, and between-host transmission (Mideo et al. 2008, Alizon et al. 2009). Existing theory and empirical studies also indicate that high virulence may be favored during the early epidemic stages, when host availability relative to inoculum pressure is high (typically at the expanding front of an epidemic), followed by a decline in virulence as the pathogen populations approach endemic equilibrium (in areas behind the front) (Berngruber et al. 2013, Griette et al. 2015, Lion & Gandon 2016, Nørgaard et al. 2021). Beyond evolutionary change, virulence is also expected to vary plastically with temperature and host resistance (*e.g*. Laine 2008, Castiblanco et al. 2020, Fidanoğlu et al. 2023, see also the reviews by Pariaud et al. 2009 and Cressler et al. 2016). Despite the central role of virulence in determining epidemic dynamics, the evolution of virulence responses to environmental variation remains largely unexplored in forest tree pathogens.

Ash dieback is an emblematic example of a emerging forest disease. It is caused by *Hymenoscyphus fraxineus*, an ascomycetous fungal pathogen native to Asia, which infects ash tree leaves during summer through wind-dispersed ascospores. These infections cause leaf wilting and necrotic lesions that can extend to the petioles and rachises, leading to premature leaf abscission (Lione et al. 2024). Lesions can also spread into and girdle the twigs and stems, damaging the cambium and inducing cankers on the bark (Gross, Holdenrieder, et al. 2014). The combined effects of leaf abscission and branch desiccation gradually lead to crown thinning and dieback, ultimately resulting in overall tree decline (Lione et al. 2024). In some regions, ash mortality rates have reached up to 85% (Coker et al. 2019). The severity of ash dieback tends to be highest under moderate summer and autumn temperatures, rather than during particularly warm periods (Grosdidier, Ioos, & Marçais 2018a, Marçais et al. 2023). Three ash species are predominantly found in Europe: the common ash (*Fraxinus excelsior* L.), the most widespread, and the narrow- leaved ash (*Fraxinus angustifolia* Vahl) and the manna ash (*Fraxinus ornus* L.), both largely restricted to southern regions. The disease particularly affects *F. excelsior* and *F. angustifolia*, whereas *F. ornus* is less susceptible (Adamčíková et al. 2018, Gross & Sieber 2016, Schwanda & Kirisits 2016).

The European epidemic of ash dieback likely results from a single introduction event, followed by the propagation of descendants of two genetically divergent strains of *H. fraxineus* (Gross, Hosoya, et al. 2014, Mcmullan et al. 2018). The disease was first reported in Poland in 1992 (Przybył 2002, Timmermann et al. 2011) and progressively reached most of the European ash distribution range (McKinney et al. 2014). Nowadays, the ash dieback epidemic has reached an endemic state in central Europe (Lygis et al. 2017, Marçais et al. 2022), but it continues to expand at the periphery of the continent, notably in southern Europe, like in Spain (Stroheker et al. 2021) and Italy (Migliorini et al. 2022). This successful invasion exemplifies the so-called genetic paradox of invasions (Estoup et al. 2016). Despite the severe initial bottleneck experienced by the fungus (Mcmullan et al. 2018), and a sharp contrast between the climatic conditions of its native and introduced ranges, *H. fraxineus* has successfully expanded across a latitudinal gradient characterized by progressively decreasing availability of susceptible hosts and climatic suitability. Indeed, as the fungus spreads southward, both temperature (Fick & Hijmans 2017) and the frequency of resistant trees increase, potentially exposing *H. fraxineus* to adaptive challenges. In this context, it is plausible that phenotypic plasticity and rapid evolutionary change in fundamental traits have facilitated the progression of the ash dieback epidemic.

To our knowledge, the evolution of virulence following *H. fraxineus* introduction into Europe has been addressed only once, by Lygis et al. (2017), who reported no difference in virulence between Lithuanian and Swiss populations, the latter located at the epidemic front at the time. However, this study did not consider the plasticity of virulence and was conducted during an earlier stage of the epidemic’s development. In *H. fraxineus*, plastic variation in virulence has so far been investigated in relation to host species (*e.g.* Schwanda and Kirisits 2016) but the evolution of plasticity in virulence during the range expansion of *H. fraxineus* remains unexplored. Furthermore, how virulence in *H. fraxineus* varies plastically with temperature has not been studied, although temperature is a key determinant of ectothermic fungal distributions (Gerken et al. 2015, McLean et al. 2005). Prior research on *H. fraxineus* has already documented thermal plasticity of *in vitro* mycelial growth (Hauptman et al. 2013, Kowalski & Bartnik 2012) and viability (Hauptman et al. 2013), with optimal in vitro growth typically observed between 22 °C and 24 °C. More recently, Bécans et al. (2025) investigated the evolution of thermal plasticity of the same traits across five European *H. fraxineus* populations sampled along a latitudinal gradient. They found that the northernmost population (Lithuania) showed reduced viability at high temperatures, whereas the southernmost population (northern Italy), located near the epidemic front, displayed significantly greater growth than other populations under elevated temperatures. As mycelial growth and survival may be related to virulence expression (Blenis et al. 1984, Kowalski & Bartnik 2012 but see Kosawang et al. 2020), these findings suggest that the thermal plasticity of virulence may be evolving during the ongoing range expansion of *H. fraxineus*, and ultimately promoting further disease spread.

In this study, we aimed to characterize the plasticity of virulence at the individual level in two distinct populations of *H. fraxineus*: one from Lithuania, where the pathogen is long-established and located near the historical center of ash dieback emergence in Europe, and another from northern Italy, representing a recently established peripheral population near the southern edge of the current epidemic front. Assuming that the southward expansion of ash dieback from its north-eastern centre of origin was continuous (Mcmullan et al. 2018), the Lithuanian population is composed of genotypes recurrently exposed to the seasonal temperature regimes characteristic of northern Europe and without prior contact with *F. ornus* (Caudullo & De Rigo 2016). In contrast, the ancestors of the Italian genotypes colonised a latitudinal gradient and were subjected both to summer heatwaves characteristic of southern Europe (Fick & Hijmans 2017) and to a host community comprising both *F. excelsior* and *F. ornus* (Caudullo & De Rigo 2016). Because expansion-front populations are typically subject to stronger genetic drift than core populations (Excoffier & Ray 2008, Excoffier et al. 2009), we first characterized neutral genetic diversity and differentiation between the two *H. fraxineus* populations as a first step toward assessing the potential contribution of genetic drift to phenotypic differences between them. We then characterized virulence reaction norms through replicated plant inoculation experiments and specifically tested the following hypotheses:

1. Uniform baseline virulence: virulence does not differ between Lithuanian and Italian *H. fraxineus* genotypes on *F. excelsior* under moderate temperature conditions.
2. Main environmental effects on virulence: virulence in pooled *H. fraxineus* genotypes decreases under an elevated temperature compared to a moderate one (a), and on *F. ornus* compared to *F. excelsior* (b).
3. Population-level differences in plasticity: Italian genotypes show lower plasticity in virulence than Lithuanian ones, with a smaller reduction in virulence when exposed to increased temperatures (a), or a more resistant *F. ornus* tree (b). This reduced plasticity is consistent with stronger phenotypic buffering in the Italian genotypes, and may lead to contrasting trait expression between the two populations under non-optimal conditions (c).

## Materials and methods

### H. fraxineus *populations*

Black, symptomatic rachises of *F. excelsior* were sampled between July 8 and the end of July 2020 at two sites, one in Lithuania and the other in Italy. Rachises were collected from the base of several contiguous symptomatic ash trees at each site. The two sites differed markedly in their distance from Estonia, where the first evidence of *H. fraxineus* presence in Europe was reported (Agan et al. 2023). These sites also differ in their altitudinal and climatic characteristics: mean annual temperature and mean summer temperature were higher at the Italian site (844 m elevation) than at the Lithuanian one (62 m). Moreover, *F. ornus* coexists with the highly susceptible *F. excelsior* in northern Italy, but is absent from Lithuania (Caudullo & De Rigo 2016) (figure 1).

**Figure 1.**
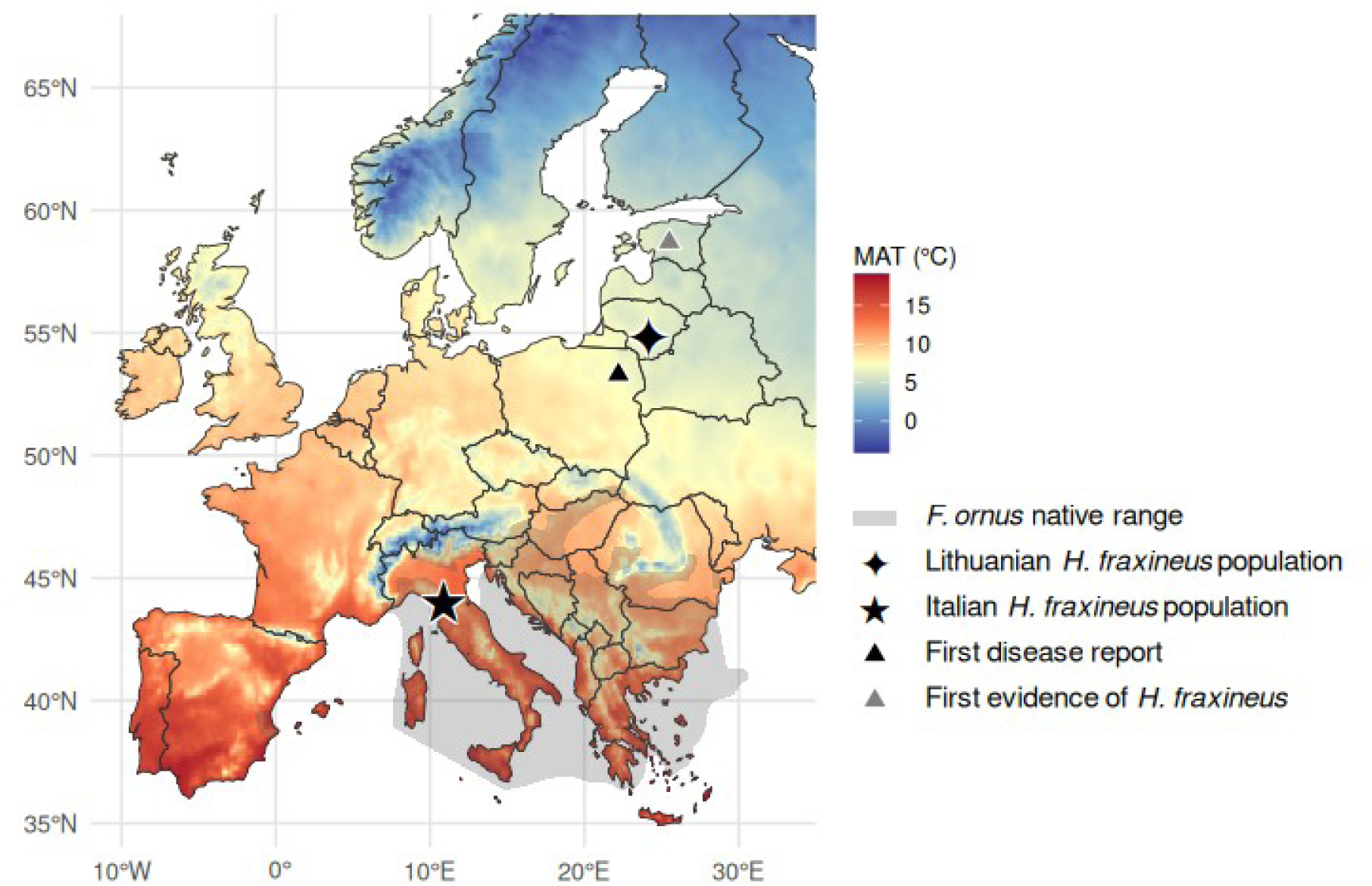
Location of the two *Hymenoscyphus fraxineus* populations studied, overlaid on mean annual temperature (MAT) and *Fraxinus ornus* native distribution (grey area) in Europe. Stars indicate the *H. fraxineus* population: Lithuania (✦, first report of ash dieback in 1996; MAT= 6.56 °C, mean summer temperature (ST) = 15.3°C) and Italy (★, first report of ash dieback in 2016; MAT = 10.05°C, ST = 17.3°C). Temperatures were obtained from WorldClim v2.1 climate normals (Fick & Hijmans, 2017) and calculated as the mean of monthly values at a spatial resolution of 18.5 km. The grey shaded area indicates the native range of *Fraxinus ornus* in Europe. Triangles mark two key events in the European ash dieback epidemic: the first evidence of *H. fraxineus* in Europe (grey triangle, Estonia) and the first reported occurrence of ash dieback (black triangle, north- eastern Poland, 1992).

Upon collection, the infected rachises were stored at 4 °C, and pathogen isolations were performed between September 17 and 19 2020, following the protocol of Kowalski & Bartnik (2012), with two modifications: ash leaf malt extract agar (AMEA, Kirisits et al. 2013) was used as the growth medium, and rachis surfaces were sterilized using 96% ethanol Fungal isolates (fungal cultures obtained from distinct rachises and grown separately) were identified as *H. fraxineus* based on culture characteristics. Species identity was confirmed by species-specific qPCR for isolates displaying atypical phenotypic profiles (see Becans et al. 2025). Based on SSR genotyping, we confirmed that all selected isolates were genetically pure and distinct from one another (see below). *H. fraxineus* isolates were maintained on AMEA at 4°C in the dark in a cold room and then incubated on a new AMEA plate three weeks before inoculation experimentations for inoculum production, at 22°C under controlled day/night cycles (16 hours light/8 hours dark). The two *H. fraxineus* populations analyzed in this study comprised the same 15 isolates (obtained from 15 rachises), as those examined by Becans et al. (2025). No formal a priori sample size calculation was performed; samples sizes were constrained by resources limitation (Lakens 2022).

### H. fraxineus DNA extraction and SSR genotyping

Mycelium of *H. fraxineus* isolates was grown for three weeks on Petri dishes covered with cellophane, then scraped and transferred into 2 mL Eppendorf tubes containing two stainless steel balls for subsequent grinding. Mycelium was lyophilized overnight prior to DNA extraction. Lysis buffer was preheated at 65°C for 10 min before adding β-mercaptoethanol (0.2% final concentration). Eight hundred microliters of buffer was added to each sample, which was vortexed and incubated at 65°C for 1 h. An equal volume (800 µL) of chloroform was added, samples were vortexed, and centrifuged for 10 min (13,000 rpm, −4°C). The aqueous phase was transferred to a fresh tube, mixed with an equal volume of isopropanol, and DNA was precipitated for 1 h at −20°C. After centrifugation (20 min), DNA pellets were washed sequentially with 70% and 95% ethanol, air-dried, and resuspended in 100 µL sterile water. DNA quantity and quality were assessed by NanoDrop spectrophotometry at the PGTB (Plateforme Génome Transcriptome de Bordeaux: https://pgtb.fr/). Genotyping was performed by the Gentyane platform (https://gentyane.clermont.inrae.fr/) using an ABI 3730 capillary sequencer (Applied Biosystems). Eighteen microsatellite (SSR) markers developed by Gross et al. (2012) were amplified, and allele sizes were scored using STRand software (Toonen & Hughes 2001).

### Plant material

Three hundred ash seedlings of *F. excelsior* and 100 ash seedlings of *F. ornus* from the Planfor nursery (Uchacq et Parentis, 40090, France), showing no visible disease symptoms, were potted in October 2022 in 4L containers and maintained outside until inoculation. On average, *F. excelsior* seedlings were 33.60 cm tall, while *F. ornus* seedlings measured 53.08 cm at the time of inoculation. Stem diameters, measured 10 cm above the soil surface, ranged from 3 to 6 mm for *F. excelsior* and from 5 to 8 mm for *F. ornus*.

### Virulence assessment of H. fraxineus isolates through inoculation of ash trees

For each of the 30 studied *H. fraxineus* isolates, we assessed virulence plasticity in response to temperature and host species separately. These assessments were based on inoculations performed under three experimental conditions: *F. excelsior* at 22°C, *F. excelsior* at 26°C, and *F. ornus* at 22°C. This design allowed us to estimate individual virulence reaction norms in response to temperature (22°C to 26°C on *F. excelsior*) and host species (*F. excelsior* to *F. ornus* at 22°C). For both types of reaction norm, *F. excelsior* at 22°C was used as the reference condition, while the treatment condition was either 26°C or *F. ornus* depending on the reaction norm considered. There is, however, considerable variation in susceptibility to ash dieback among common ash trees, which has been associated with differences in iridoid glycoside content (Sollars et al. 2017). Because 30 *H. fraxineus* isolates could not be inoculated on the same ash tree, variation in host susceptibility to *H. fraxineus* may have biased our assessment of population-level differences in pathogen virulence and its plasticity. Previous work has shown that analyses of phenolic bark extracts using Fourier- transform infrared spectrometry, can successfully discriminate among ash trees according to their susceptibility to *H. fraxineus* (Villari et al. 2018). To limit intraspecific variation in resistance among the *F. excelsior* (susceptible) and *F. ornus* (resistant) seedlings used in this study, we implemented the following strategy. We first analyzed the near-infrared spectra of the seedlings using an approach adapted from Villari et al. (2018), relying on leaf-level near-infrared spectrometry and principal component analysis of the spectral data. The first principal component (PC1, 86.3% of the total variance) mainly captured variation among *F. excelsior* seedlings, while the two ash species segregated along PC2 (9.0%) (figure S1). Three *F. ornus* seedlings overlapped with the *F. excelsior* cluster. These three individuals, together with those located at the margins of their species cluster, were excluded from the inoculation experiments. The retained susceptible group comprised 46 *F. excelsior* seedlings, while the resistant group included 23 *F. ornus* seedlings. To further minimize the effects of residual variation in host resistance on the expression of virulence, each isolate was inoculated on three distinct trees under each experimental condition ( *F. excelsior* at 22°C, *F. excelsior* at 26°C, and *F. ornus* at 22°C). Each seedling was inoculated with a unique combination of four isolates, consisting of two Italian and two Lithuanian isolates, with the exception of two seedlings, which received three isolates from one population and one from the other. In addition, each tree received a mock inoculation with a sterile AMEA plug. Foliar inoculations were carried out from July 19 to 21 2023, following the protocol by Orton et al. (2019), except that we used standardized AMEA plugs colonized by *H. fraxineus* mycelium instead of mycelium agar mix. A 5 mm AMEA plug of three-week-old mycelium was placed on the epidermal tissues of a rachis between the second and third pair of leaflets, at a site that had been preliminary wounded with a standardized 5 mm incision. The inoculum was maintained in place and prevented from drying by wrapping the inoculation site with parafilm, which was removed seven days after inoculation. The plants were maintained in separate climate chambers set at 22°C or 26°C, with controlled day/night cycles (16 h light / 8 h dark) and a constant relative humidity of 70%. In total, 345 foliar inoculations were performed on 69 ash seedlings: 23 *F. excelsior* seedlings exposed to 22 °C, 23 *F. excelsior* seedlings exposed to 26 °C, 23 *F. ornus* seedlings exposed to 22 °C, with three replicate inoculations per *H. fraxineus* isolate on distinct seedlings for each experimental condition. To complete the inoculation scheme across all available ash seedlings, six randomly selected *H. fraxineus* isolates were inoculated in additional replicates. These additional replicates were not included in subsequent analyses, which were restricted to a maximum of three replicates per isolate. Prior to inoculation, the isolates were anonymized to prevent potential bias during experimental monitoring. We assessed the virulence using three complementary proxies that are associated with distinct stages of infection:

i. lesion length, a continuous variable representing the extent of the necrosis (in cm), which informs about pathogen colonization.
ii. symptom occurrence, a binary variable indicating whether pathogen establishment generated symptoms on the inoculated leaf (0: no symptoms 1: symptoms present),
iii. symptom severity, a binary variable recorded conditionally only for symptomatic leaves, indicating whether the symptoms became severe (0: no severe wilting 1: severe wilting or leaf shedding).

We monitored lesion length and leaf symptoms weekly, starting two weeks after inoculation and continuing until the ninth week. We stopped measuring lesions at week seven because they could no longer be reliably distinguished after leaf fall. Throughout the manuscript, “symptom severity” refers to symptom progression to a severe state.

### Population genetic analysis

We performed genetic analyses in R using the packages poppr (v. 2.9.8), ade4 (v. 1.7-24), and hierfstat (v. 0.5.11). Multilocus genotypes were defined as unique combinations of alleles across the 18 SSR loci. We quantified genotypic diversity using the number of unique multilocus genotypes, the Shannon-Wiener diversity index, Simpson’s index, genotypic evenness, and the index of association to assess multilocus linkage disequilibrium. We assessed genetic differentiation between populations using an analysis of molecular variance and principal component analysis based on allele frequencies. The significance of population differentiation was assessed using 999 permutations. Finally, we estimated per-locus F*ST* values using hierfstat (v. 0.5.11).

### Statistical analysis of reaction norms diversity

For each experiment, we analyzed virulence trait variation across the two pathogen populations, the two environments and time, using generalized linear mixed-effects models (GLMMs) fitted with the glmmTMB package (v. 1.1.4) in R (v. 4.3.3, R Core Team 2021). We modeled Lesion length (continuous quantitative) using Tweedie distributions, and modeled symptom onset and severe symptoms (both binary) using binomial distributions.

Full initial models included the fixed effects of *Population*, *Treatment* (temperature: 22°C *vs* 26°C, or host species: *F. excelsior vs F. ornus*), Days after inoculation (*Day*), and all their interactions. Non-linear temporal dynamics were modeled by applying natural cubic splines with three degrees of freedom to *Day*. Models included random intercepts for host seedlings (*u*), *H. fraxineus* isolates (*v*), and leaves (*w*), the latter accounting for the non-independence of repeated measurements from the same lesion over time. A compact description of the model is:

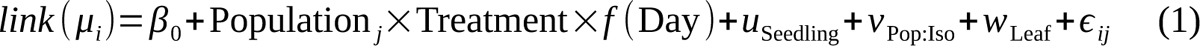

where *μ_i_* is the expected value of the response variable for observation *i* (lesion length (cm), or probability of symptom onset / severe symptoms, depending on the model), *β*_0_ is the intercept (overall estimated mean on the link scale) and ε the residual variation. The *link* function corresponds to the distribution used: log link for lesion length (Tweedie), logit link for the binary traits (binomial).

We performed model simplification for fixed effects from Akaike Information Criterion (AIC) and Likelihood Ratio Tests (LRTs, χ^2^-test, α = 0.05), removing three-way interactions when their removal did not result in a significant loss of model fit. We simplified random-effect structures, when necessary, to resolve model assumptions violation, convergence issues or singular fits. Model assumptions and residual diagnostics (*e.g*. dispersion, zero-inflation, temporal autocorrelation) were systematically inspected using the DHARMa package (v0.5.0).

We assessed the statistical significance of fixed effects and their interactions from estimated marginal means and associated 95% confidence intervals, computed on the *link* scale using the emmeans package (v. 2.0.3). We computed post-hoc pairwise contrasts between *H. fraxineus* populations across monitoring dates using estimated marginal means (*z*-score, *α* = 0.05). These contrasts were calculated for each virulence component under reference conditions (22 °C and *F. Excelsior,* hypothesis 1), under treatment conditions (26 °C or *F. ornus*) (hypotheses 3a and 3b), and to assess phenotypic plasticity, defined as the difference in trait values across conditions (hypotheses 3a and 3b). We systematically adjusted *p*-values for multiple testing across dates using the conservative Bonferroni correction (*α* = 0.05). Following the same pairwise contrast procedure, we further evaluated the overall effects of *Day* or *Treatment* (hypotheses 2a and 2b) on the virulence components and their plasticity in the pooled *H. fraxineus* genotypes. For each monitoring day, we estimated point-biserial correlations and their significancy between lesion length and symptom occurrence, and lesion length and the probability of symptom becoming severe (stats package, v. 4.3.3, *α* = 0.05). Data manipulation and figures construction were carried out using dplyr (v. 1.2.1), tidyr (v. 1.3.2), ggplot2 (v. 4.0.3).

## Results

### The two H. fraxineus populations show high genotypic diversity but limited genetic differentiation

All 30 isolates possessed distinct multilocus genotypes. Indices of association were close to zero in both populations, indicating no detectable multilocus linkage disequilibrium and supporting predominantly recombining rather than clonal population structures (table S1).

Principal component analysis (with PC1 = 13.8% and PC2 = 11.7% of the total variance) did not show any clear population-level separation (figure S2). According to the analysis of molecular variance, most molecular variance occurred within populations (95.1%), whereas only 4.9% was attributable to differences between populations (*Φ* = 0.049, *p* = 0.034). Per-locus FST values were close to zero for most SSR markers, with only two loci exceeding 0.03 (maximum = 0.046, figure S3), supporting weak genetic differentiation between the Italian and Lithuanian populations.

### Overall dynamics of virulence expression in the inoculation experiment

Throughout the experiment, both the estimated mean lesion length and the probability of severe symptoms increased significantly over the monitoring period (figure 2a, g, table 1). In contrast, the estimated mean probability of symptom onset in inoculated rachises was already high early in the experiment and remained stable thereafter. For instance, pooled *H. fraxineus* isolates induced symptoms in *F. excelsior* at 22°C with an estimated probability of 0.86 (95% confidence interval [0.67, 0.95]) at day 14 and 0.91 [0.75, 0.97] at day 63 (figure 2d). In contrast, under the same conditions, the estimated probability of symptom progression to a severe state increased significantly from 0.05 [0.00, 0.25] at day 14 to 0.98 [0.94 1.00] at day 63 (figure 2g). No lesions or symptoms developed following control inoculations, and rachis lesions never extended onto seedling trunks during the monitoring period.

**Figure 2.**
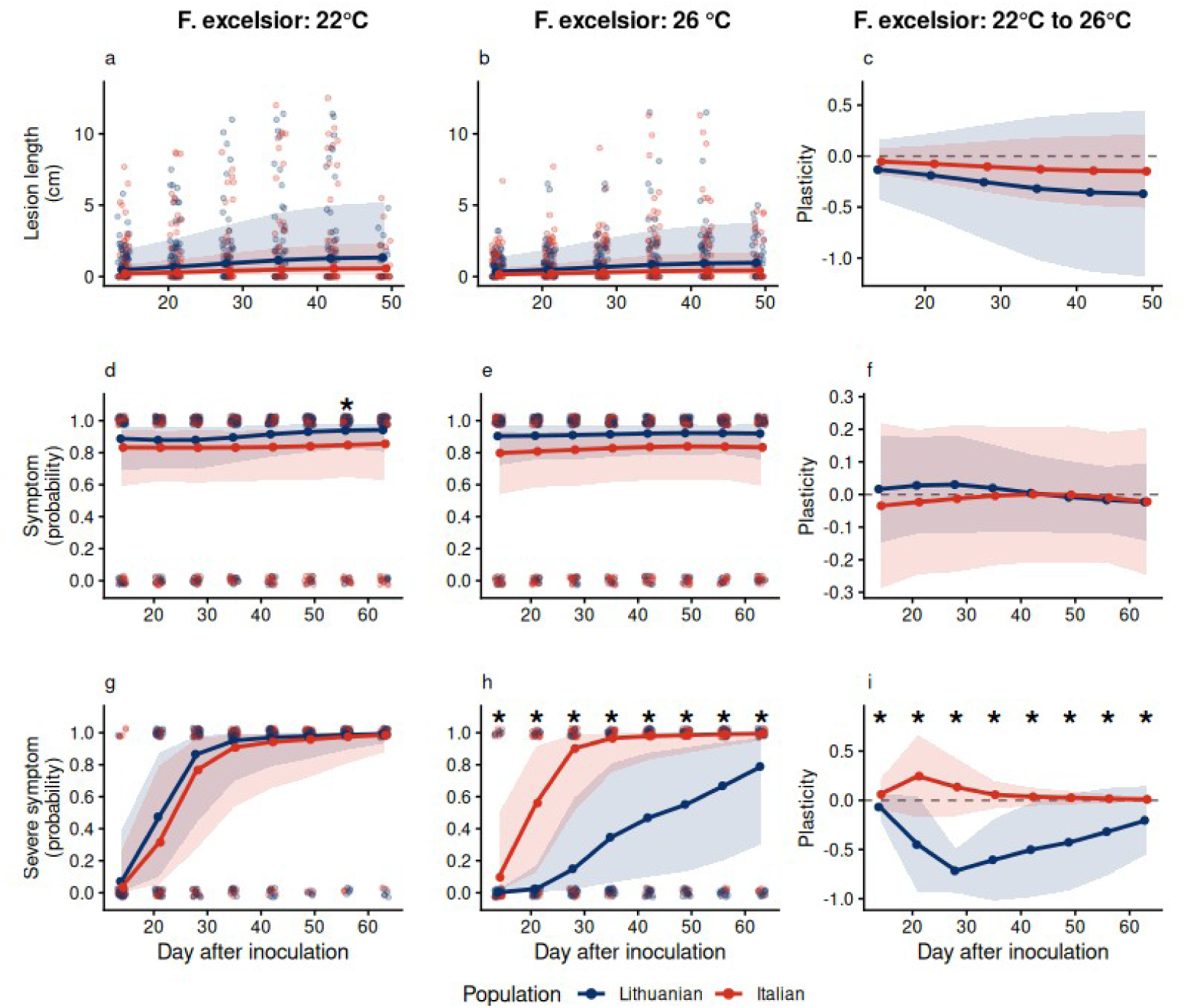
Estimated trait values and temperature-induced plasticity for lesion length (a, b, c), symptom probability (d, e, f), and severe symptom probability (g, h, i), from day 14 to day 63 post- inoculation, in two *Hymenoscyphus fraxineus* populations. Columns show the estimated trait values under the reference condition (*Fraxinus excelsior*, 22°C) (a, d, g), under elevated temperature (*Fraxinus excelsior*, 26°C) (b, e, h), and the estimated slopes of the reaction norms (c, f, i). Bold points represent marginal means for each population, light-colored points indicate isolate-level means. Shaded areas represent 95% confidence intervals. Stars indicate significant pairwise differences between populations (α = 0.05).

**Table 1.** Summary of post-hoc pairwise comparisons of estimated marginal means. Pairwise comparisons were performed to assess differences in lesion length, symptom occurrence, and symptom severity between sampling days, temperatures, ash species, and *H. fraxineus* populations. The *Effect* column indicates the factor being compared, *Condition* specifies the environmental condition(s) under which the comparisons were performed, *Contrast* indicates the levels being compared and *Trait* indicates the virulence component considered. *Significant period* indicates the sampling days over which a significant difference was detected. *z-scores* and adjusted *p*-values are reported as ranges when their values varied among comparisons.

| Effect | Condition | Contrast | Trait | Significant period | z-score | Adjusted-p |
|---|---|---|---|---|---|---|
| Day | *F. ex.*/22°C | Pairwise consecutive days | Lesion length | Day 14 to 42 | <-5.39 | **<0.01** |
|  | *F. ex.*/26°C | Pairwise consecutive days | Lesion length | Day 14 to 42 | <-5.39 | **<0.01** |
|  | *F. or.*/22°C | Pairwise consecutive days | Lesion length | Day 21 to 35 | <-4.12 | **<0.01** |
|  | *F. ex.*/22°C | Pairwise consecutive days | Symptom | None | -1.27 to 0.21 | >0.20 |
|  | *F. ex.*/26°C | Pairwise consecutive days | Symptom | None | -0.65 to 0.15 | >0.52 |
|  | *F. or.*/22°C | Pairwise consecutive days | Symptom | None | -0.94 to 0.15 | >0.35 |
|  | *F. ex.*/22°C | Pairwise consecutive days | Severe symptom | Day 14 to 42 | -7.94 to -1.33 | **<0.04** |
|  | *F. ex.*/26°C | Pairwise consecutive days | Severe symptom | Day 14 to 63 | -9.52 to -2.18 | **<0.03** |
|  | *F. or.*/22°C | Pairwise consecutive days | Severe symptom | Day 21 to 63 | -5.64 to -1.40 | **<0.02** |
| Temperature | *F. ex.*/22°C, *F. ex.*/26°C | *F. ex.*/22°C, *F. ex.*/26°C | Lesion length | None | 1.43 | 0.92 |
|  | *F. ex.*/22°C, *F. ex.*/26°C | *F. ex.*/22°C, *F. ex.*/26°C | Symptom | None | -0.16 to 0.32 | >0.75 |
|  | *F. ex.*/22°C, *F. ex.*/26°C | *F. ex.*/22°C, *F. ex.*/26°C | Severe symptom | None | 1.52 | 1.00 |
| Host | *F. or.*/22°C, *F. or.*/22°C | *F. or.*/22°C, *F. or.*/22°C | Lesion length | Day 14 to 49 | 3.85 | **<0.01** |
|  | *F. or.*/22°C, *F. or.*/22°C | *F. or.*/22°C, *F. or.*/22°C | Symptom | Day 14 to 63 | 5.80 | **<0.01** |
|  | *F. or.*/22°C, *F. or.*/22°C | *F. or.*/22°C, *F. or.*/22°C | Severe symptom | Day 14 to 63 | 3.35 to 7.26 | **<0.01** |
| Population | *F. ex.*/22°C | Lithuania, Italy | Lesion length | None | 0.85 | 1.00 |
|  | *F. ex.*/22°C | Lithuania, Italy | Symptom | Day 56 | 2.80 | **0.04** |
|  | *F. ex.*/22°C | Lithuania, Italy | Severe symptom | None | 0.53 | 1.00 |
|  | *F. ex.*/26°C | Lithuania, Italy | Lesion length | None | 0.82 | 0.41 |
|  | *F. ex.*/26°C | Lithuania, Italy | Symptom | None | 1.50 to 2.48 | >0.11 |
|  | *F. ex.*/26°C | Lithuania, Italy | Severe symptom | Day 14 to 63 | -2.99 | **0.02** |
|  | *F. ex.*/22°C to *F. ex.*/26°C | Lithuania, Italy | Lesion length | None | 0.07 | 0.95 |
|  | *F. ex.*/22°C to *F. ex.*/26°C | Lithuania, Italy | Symptom | None | -0.96 to 0.37 | 1.00 |
|  | *F. ex.*/22°C to *F. ex.*/26°C | Lithuania, Italy | Severe symptom | Day 14 to 63 | 6.43 | **<0.01** |
|  | *F. or.*/22°C | Lithuania, Italy | Lesion length | None | -0.02 | 1.00 |
|  |  | Lithuania, Italy | Symptom | None | 0.53 | 1.00 |
|  |  | Lithuania, Italy | Severe symptom | None | -1.78 to -0.04 | >0.08 |
|  | F. ex./22°C to F. or./22°C | Lithuania, Italy | Lesion length | None | 2.61 | >0.05 |
|  |  | Lithuania, Italy | Symptom | None | -0.58 | 1.00 |
|  |  | Lithuania, Italy | Severe symptom | None | -0.20 to 2.55 | >0.09 |

Between days 14 and 49, lesion length was strongly correlated with symptom occurrence ( *r* = 0.49- 0.81, *p* < 0.01). A significant correlation was also observed between lesion length and the probability of severe symptoms (*r* = 0.35-0.71, *p* < 0.01) from day 21 onwards, although the strength and timing of this association varied among days (table S2).

### Lithuanian and Italian H. fraxineus isolates expressed similar baseline virulence in the reference environment (hypothesis 1)

The two *H. fraxineus* populations caused lesions of similar length and symptoms of similar severity on *F. excelsior* rachises at 22 °C (figure 2a, g table 1). The probability of symptom induction was also generally similar between the two *H. fraxineus* populations, except at day 56, when Lithuanian isolates induced symptoms more frequently than Italian isolates (0.94 *vs* 0.85, *z* = 2.80, *p* = 0.04, figure 2d). Note however that variation in lesion length and symptom induction was primarily associated with differences among isolates and seedlings rather than with population origin or other fixed effects. Indeed, fixed effects explained little variation in these traits (marginal *R*^2^, *R*^2^m ≤ 0.04), whereas including random effects substantially increased the variance explained (conditional *R*^2^, *R*^2^c ≥ 0.61; tables 2 and 3). In contrast, fixed effects explained more than 29% of the variation in symptom severity, while the full model explained 89%.

**Table 2.** Summary of the generalized linear mixed models testing the effect of temperature on the virulence components (lesion length, symptom, symptom severity). *SE*: standard error, *SD*: standard deviation, *z* and *p*: *z*-ratio and *p*-value from Wald tests, *R²m* and *R²c*: marginal and conditional R^2^.

| Trait | R<sup>2</sup>m | R<sup>2</sup>c | Fixed effect |  |  |  |  | Random effect |  |
|---|---|---|---|---|---|---|---|---|---|
|  |  |  | Variable | Estimate | SE | *z* | *p* | Variable | SD |
| Lesion length | 0.04 | 0.99 | Population | -0.84 | 0.99 | -0.85 | 0.40 | Lesion | 1.14 |
|  |  |  | Temperature | -0.32 | 0.29 | -1.13 | 0.26 | Tree | 0.33 |
|  |  |  | ns(Day)1 | 0.83 | 0.06 | 14.35 | **<0.01** | Population: Isolate | 2.55 |
|  |  |  | ns(Day)2 | 1.39 | 0.09 | 15.48 | **<0.01** |  |  |
|  |  |  | ns(Day)3 | 0.76 | 0.05 | 16.31 | **<0.01** |  |  |
|  |  |  | Population*Temperature | 0.03 | 0.39 | 0.07 | 0.95 |  |  |
| Symptom | 0.02 | 0.61 | Population | -0.46 | 0.52 | -0.88 | 0.38 | Tree | 2.24 |
|  |  |  | Temperature | 0.18 | 0.89 | 0.20 | 0.84 |  |  |
|  |  |  | ns(Day)1 | 0.66 | 0.68 | 0.98 | 0.33 |  |  |
|  |  |  | ns(Day)2 | 0.51 | 0.98 | 0.52 | 0.60 |  |  |
|  |  |  | ns(Day)3 | 0.87 | 0.50 | 1.73 | 0.08 |  |  |
|  |  |  | Population*Temperature | -0.41 | 0.74 | -0.55 | 0.58 |  |  |
|  |  |  | Population*ns(Day)1 | -0.63 | 0.90 | -0.70 | 0.48 |  |  |
|  |  |  | Population*ns(Day)2 | -0.39 | 1.34 | -0.297 | 0.77 |  |  |
|  |  |  | Population*ns(Day)3 | -0.68 | 0.65 | -1.038 | 0.30 |  |  |
|  |  |  | Temperature*ns(Day)1 | -0.39 | 0.95 | -0.40 | 0..69 |  |  |
|  |  |  | Temperature*ns(Day)2 | -0.23 | 1.42 | -0.17 | 0.87 |  |  |
|  |  |  | Temperature*ns(Day)3 | -0.71 | 0.69 | -1.03 | 0.30 |  |  |
|  |  |  | Population*Temperature*ns(Day)1 | 0.64 | 1.27 | 0.51 | 0.61 |  |  |
|  |  |  | Population*Temperature*ns(Day)2 | 0.51 | 1.92 | 0.26 | 0.79 |  |  |
|  |  |  | Population*Temperature*ns(Day)3 | 0.67 | 0.91 | 0.74 | 0.46 |  |  |
| Severe symptom | 0.29 | 0.89 | Population | -0.67 | 1.28 | -0.52 | 0.60 | Tree | 2.80 |
|  |  |  | Temperature | -3.60 | 1.28 | -0.53- | **<0.01** | Population: Isolate | 3.07 |
|  |  |  | ns(Day)1 | 4.60 | 0.58 | 7.88 | **<0.01** |  |  |
|  |  |  | ns(Day)2 | 12.00 | 1.15 | 10.42 | **<0.01** |  |  |
|  |  |  | ns(Day)3 | 3.98 | 0.47 | 8.46 | **<0.01** |  |  |
|  |  |  | Population*Temperature | 4.62 | 0.72 | 6.42 | **<0.01** |  |  |

**Table 3.** Summary of the generalized linear mixed models testing the effect of Host (*Fraxinus excelsior*: susceptible, *Fraxinus ornus*: resistant) on the virulence components (lesion length, symptom, symptom severity). *SE*: standard error, *SD*: standard deviation, *z* and *p*: *z*-ratio and *p*- value from Wald tests, *R²m* and *R²c*: marginal and conditional *R*^2^.

| Trait | R<sup>2</sup>m | R<sup>2</sup>c | Fixed effect |  |  |  |  | Random effect |  |
|---|---|---|---|---|---|---|---|---|---|
|  |  |  | Variable | Estimate | SE | *z* | *p* | Variable | SD |
| Lesion length | 0.07 | 0.94 | Population | -0.30 | 0.61 | -0.50 | 0.62 | Tree | 0.72 |
|  |  |  | Host | -1.02 | 0.23 | -4.39 | **<0.01** | Population: Isolate | 1.63 |
|  |  |  | ns(Day)1 | 0.73 | 0.11 | 6.49 | **<0.01** |  |  |
|  |  |  | ns(Day)2 | 1.09 | 0.17 | 6.31 | **<0.01** |  |  |
|  |  |  | ns(Day)3 | 0.57 | 0.08 | 6.75 | **<0.01** |  |  |
|  |  |  | Population*Host | 0.32 | 0.12 | 2.61 | **<0.01** |  |  |
| Symptom | 0.02 | 0.75 | Population | -0.45 | 1.21 | -0.37 | 0.71 | Isolate | 3.10 |
|  |  |  | Host | -0.87 | 0.23 | -3.71 | **<0.01** |  |  |
|  |  |  | ns(Day)1 | 0.31 | 0.35 | 0.90 | 0.37 |  |  |
|  |  |  | ns(Day)2 | 0.52 | 0.52 | 1.00 | 0.32 |  |  |
|  |  |  | ns(Day)3 | 0.33 | 0.25 | 1.32 | 0.19 |  |  |
|  |  |  | Population*Host | -0.19 | 0.33 | -0.58 | 0.56 |  |  |
| Severe symptom | 0.56 | 0.89 | Population | 0.64 | 1.31 | 0.49 | 0.62 | Tree | 2.14 |
|  |  |  | Host | -5.17 | 2.04 | -2.54 | **0.01** | Isolate | 2.36 |
|  |  |  | ns(Day)1 | 4.33 | 1.02 | 4.25 | **<0.01** |  |  |
|  |  |  | ns(Day)2 | 13.79 | 2.03 | 6.81 | **<0.01** |  |  |
|  |  |  | ns(Day)3 | 3.35 | 0.80 | 4.18 | **<0.01** |  |  |
|  |  |  | Population*Host | -0.54 | 2.66 | -0.20 | 0.84 |  |  |
|  |  |  | Population*ns(Day)1 | 0.91 | 1.56 | 0.58 | 0.56 |  |  |
|  |  |  | Population*ns(Day)2 | -2.25 | 2.55 | -0.88 | 0.38 |  |  |
|  |  |  | Population*ns(Day)3 | 1.19 | 1.31 | 0.91 | 0.36 |  |  |
|  |  |  | Host*ns(Day)1 | 1.71 | 1.99 | 0.86 | 0.39 |  |  |
|  |  |  | Host*ns(Day)2 | -5.22 | 4.15 | -1.26 | 0.21 |  |  |
|  |  |  | Host*ns(Day)3 | 4.80 | 1.56 | 3.09 | **<0.01** |  |  |
|  |  |  | Population*Host*ns(Day)1 | -0.52 | 2.69 | -0.20 | 0.85 |  |  |
|  |  |  | Population*Host*ns(Day)2 | 6.53 | 5.99 | -1.09 | 0.28 |  |  |
|  |  |  | Population*Host*ns(Day)3 | -1.64 | 2.03 | -0.81 | 0.42 |  |  |

### Overall H. fraxineus virulence was reduced on F. ornus, but was not significantly affected by temperature (hypotheses 2a and 2b)

When pooling all *H. fraxineus* isolates, temperature had no significant effect on lesion length, symptom development, or symptom severity over the monitoring period (hypothesis 2a, figure 2 panels a, b, d, e, table 1). In contrast, all three components of virulence were significantly lower on *F. ornus* than on *F. excelsior* throughout the experiment (z ≥ 3.35, p < 0.01) (hypothesis 2b, figure 3, panels a, b, d, e, table 1). For example, on day 35, the probability of symptom development in inoculated rachises was estimated at 0.88 [0.68, 0.96] on *F. excelsior*, compared with 0.74 [0.45, 0.90] on *F. ornus*. Among symptomatic rachises, the probability of progression to severe state was 0.93 [0.75, 0.98] on *F. excelsior*, compared with less than 0.01 [0.00, 0.04] on *F. ornus*. However, this difference decreased over time, with severe symptom probability reaching 0.99 [0.93 1.00] on *F. excelsior* and 0.64 [0.28, 0.89] on *F. ornus* by day 63.

**Figure 3.**
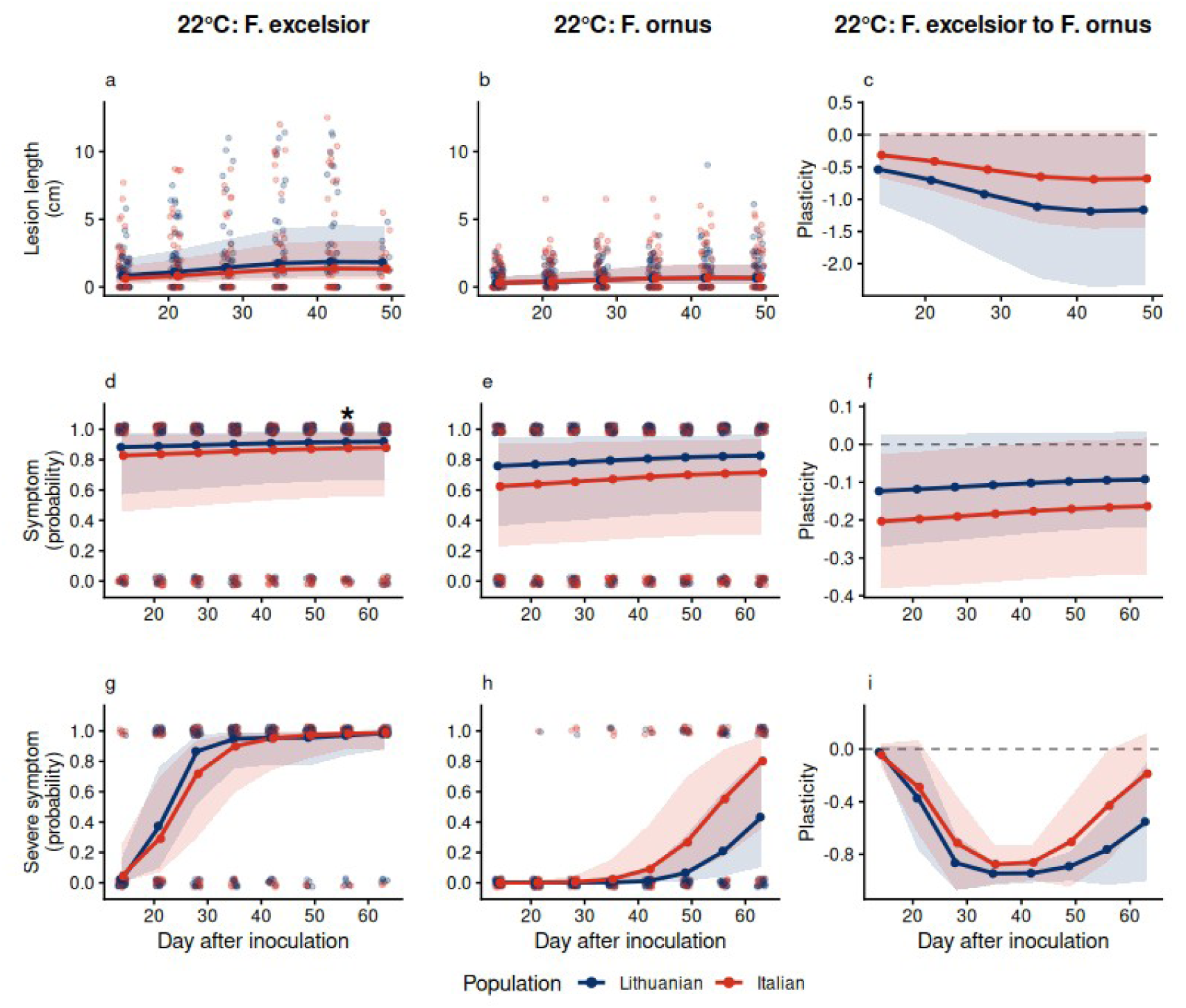
Estimated trait values and host-induced plasticity for lesion length (a, b, c), symptom probability (d, e, f), and severe symptom probability (g, h, i), from day 14 to day 63 post- inoculation, in two *Hymenoscyphus fraxineus* populations. Columns show the estimated trait values under the reference condition (*Fraxinus excelsior*, 22°C) (a, d, g), on *Fraxinus ornus* (22°C) (b, e, h), and the estimated slopes of the reaction norms (c, f, i). Bold points represent marginal means for each population, light-colored points indicate isolate-level means. Shaded areas represent 95% confidence intervals. Stars indicate significant pairwise differences between populations (α = 0.05).

### Italian isolates caused more severe symptoms than Lithuanian isolates under elevated temperatures (hypotheses 3a and 3c)

The thermal plasticity of lesion length and symptom induction did not differ between the *H. fraxineus* populations throughout the experiment (figure 2c, f, table 1). Likewise, at 26°C, neither lesion length nor symptom induction differ between populations (figure 2b, e, table 1).

In contrast, severe symptom induction was less negatively impacted by temperature increase in Italian than in Lithuanian isolates throughout the experiment (*z* = 6.42, *p* < 0.01, figure 2i). The estimated reaction-norm slope for the Italian population did not differ from zero at any time point, indicating no significant thermal plasticity in symptom severity. In contrast, the Lithuanian reaction norm showed a consistently negative slope from day 14 to 63 (*e.g.* −3.60/°C on day 35 [−5.43, −1.77], figure 2i). At 26°C, symptom severity was also higher in seedlings inoculated with Italian isolates than in those inoculated with Lithuanian isolates, at all monitored dates (z = −2.99, *p* = 0.02, figure 2h). For instance, at day 35, the probability of severe symptom induction reached 0.96 [0.75 1.00] in Italian isolates, versus 0.35 [0.06, 0.81] in Lithuanian isolates.

### Similar virulence plasticity in relation to ash species between Lithuanian and Italian H. fraxineus populations (hypothesis 3b)

Across the monitoring period, the estimated reaction norms for the three virulence components as a function of ash species were similar between the two *H. fraxineus* populations (figure 3 panels c, f, i, table 1). Depending on the monitoring day, the estimated reaction norms’ slopes were either negative or not significant from zero in both populations. At 35 days post-inoculation, reaction norm slopes for lesion length and symptom induction were −0.70 cm/°C [−1.15, 0.25] and −1.06/°C [−1.52, 0.60] in the Italian population, and −1.02 cm/°C [1.47, −0.56] and −0.87/°C [−1.32, 0.41] in the Lithuanian population, respectively. At this time point, the probability of severe symptoms in symptomatic *F. ornus* seedlings remained close to zero in both populations, but subsequently increased, reaching 0.43 [0.10, 0.83] in Lithuanian and 0.80 [0.36, 0.88] in Italian isolates on the last day of monitoring. Italian isolates thus tended to cause more severe symptoms than Lithuanian ones at the end of the monitoring period, although this difference did not reach significance (figure 3 panels h and i, *p* > 0.08).

## Discussion

We characterized reaction norms for virulence in Lithuanian and Italian populations of *H. fraxineus*, an invasive fungal pathogen responsible for the ash dieback epidemic currently affecting Europe. The Lithuanian population was sampled near the presumed site of the initial emergence of the disease, whereas the Italian population is more recently established and located near the southern edge of the epidemic front. The two studied *H. fraxineus* populations showed weak genetic differentiation and similarly high levels of genotypic diversity. Using leaf inoculations, we assessed variation in the thermal and host-related plasticity of three virulence proxies: lesion length, symptom induction, and symptom severity. Under reference conditions (*F. excelsior* 22 °C), virulence was comparable between the two *H. fraxineus* populations. Virulence was reduced on *F. ornus* relative to *F. excelsior* in both populations, with similar responses between populations for all three traits. Increasing temperature had no effect on lesion length or symptom induction in either population. However, we found that the probability of severe symptom induction decreased with increasing temperature in the Lithuanian population but remained stable across temperatures in the Italian population, indicating canalization of this trait. In other words, once established on *F. excelsior*, the Italian genotypes were more harmful to the host at high temperatures than the ancestral Lithuanian genotypes.

### H. fraxineus may cause high mortality rates in infected ashes despite high temperatures in souhern Europe

Among the three proxies of virulence we assessed, only the occurrence of severe foliar symptoms (pronounced wilting or premature abscission) allowed us to discriminate between the two *H. fraxineus* populations, even though lesion length was substantially correlated with both symptom induction and severe symptom induction. From a methodological point of view, this result supports the use of multiple, closely related proxies at biologically relevant scale when quantifying variation in virulence (Bock et al. 2022), as recommended for *H. fraxineus* by Orton et al. (2019). Ash dieback severity has often been linked to the capacity of *H. fraxineus* to extend infections from leaves into woody tissues, but this ability has also been regarded as an epidemiological dead end for the fungus because stem infections do not contribute to apothecia formation and onward transmission (Gross et al. 2014, Marçais et al. 2023). Conversely, accelerated leaf abscission induced by foliar infections facilitate a more rapid transition into the saprotrophic phase in the litter, thereby reducing exposure to the high canopy temperatures typical of sunny periods of southern Europe, and potentially improving strain viability (Hauptman et al. 2013, Becans et al. 2025). Severe and recurrent defoliation caused by *H. fraxineus* weakens infected ashes (Combes et al. 2024), increasing their susceptibility to secondary fungal pathogens that further accelerate decline (Chandelier et al. 2016, Benigno et al. 2023, Peters et al. 2023, Spiegel et al. 2025). Necrotic leaf lesions can also impair photosynthetic efficiency and reduce carbohydrate storage, which in turn further constrains photosynthesis, radial growth and hydraulic function, thereby increasing the likelihood of mortality in infected trees (Klesse et al. 2021a). Consequently, the evolution towards reduced thermal plasticity of virulence in southern Europe could enable the fungus to cause damages on *F. excelsior* comparable to those already documented in central Europe, despite high summer temperature conditions. Note however that future epidemic severity in this area will also be influenced by other key factors, such as host density (Martin et al. 2025) and humidity level (Sturrock et al. 2011, Klesse et al. 2021b).

### Limited initial genetic diversity in the founding population may have constrained the evolution of baseline virulence in H. fraxineus

Lygis et al. (2017) investigated the evolution of *H. fraxineus* virulence in Europe and reported high variability among genotypes (also reported by Orton et al. (2019) and Kosawang et al.(2020)), but no difference in virulence between a long-established Lithuanian population and a recently established Swiss population, the latter located at the epidemic front at the time. Although our analysis was based on a relatively small number of genotypes and only two populations, the absence of differentiation in baseline virulence between the Italian and Lithuanian *H. fraxineus* populations under reference conditions (*F. excelsior*, 22 °C) is consistent with these findings.

Several models predict that competition among pathogen genotypes for host resources increases virulence, especially when individual hosts are coinfected by multiple genotypes or species (Bashey 2015, Tollenaere et al. 2016), a common feature of ash dieback. Other model predicts that spatially varying relationships between host availability and mean virulence can drive spatial divergence in virulence between the epidemic front and area behind the front (*e.g.* Lion and Gandon 2016). Although these predictions have received empirical support in various pathosystems (*e.g*., Fenner & Ratcliffe 1965, Tack et al. 2012, Berngruber et al. 2013, Fraser et al. 2014), they do not seem to align with the findings of Lygis et al. (2017) or with our results.

The strong genetic bottleneck undergone by *H. fraxineus* during its introduction in Europe (Mc Mullan et al. 2018) likely coincided with a decrease in its virulence against *F. excelsior*, in comparison with genotypes from the native range of the fungus in Japan (Gross & Sieber 2016). Interestingly, our results suggest that baseline virulence subsequently remained stable during the spread of *H. fraxineus* towards southern Europe. One possible interpretation is that the weak initial genetic variation in the European *H. fraxineus* founding population together with founder effects and genetic drift may have limited the fixation of new alleles allowing for increased virulence expression. Nonetheless, the high genotypic diversity observed within populations coupled to the weak genetic differentiation observed between the Italian and Lithuanian populations provide little evidence for strong cumulative founder effects during the spread of the fungus. It is also possible that the extensive gene flow occurring among European *H. fraxineus* populations (Burokiene et al. 2015; Grosdidier, et al. 2018b; Gross, Hosoya, et al. 2014) together with the recent establishment of the pathogen in the Italian peninsula (Luchi et al. 2016) may have limited differentiation in baseline virulence during epidemic progression.

In addition, the coexistence of susceptible and resistant hosts could theoretically have favored increased baseline virulence, particularly if resistance in *F. ornus* imposed directional selection on pathogen virulence (Gandon & Michalakis 2000, see also Soularue et al. 2025). Yet, the low number of *H. fraxineus* genotypes infecting *F. ornus* trees may have limited the contribution of this host to virulence evolution at the regional scale in Italy (Kirisits 2017). This overall suggests that the biotic and abiotic conditions typical of southern Europe may not have been conducive to further evolution of baseline virulence.

### Differentiation in thermal reaction norm of virulence suggests thermal adaptation in H. fraxineus at the southern epidemic front

This is the second study to report phenotypic differentiation among European *H. fraxineus* populations for a trait related to thermal performance. Bécans et al. (2025) found that Italian isolates had a growth rate at 26 °C twice as high as that of Lithuanian isolates, as well as greater viability at temperatures above 30 °C. In our study, we found differences in thermal virulence plasticity between these same two populations, which are consistent with the *in vitro* results obtained by Bécans et al. (2025). Taken together, these results suggest that variation in *in vitro* thermal performance may be associated with variation in pathogen virulence. This interpretation is consistent with the hypothesis proposed by Blenis et al. (1984) and with previous *H. fraxineus* leaf inoculation experiments (Orton et al. 2019; but see Kosawang et al. 2020).

The Italian genotypes and their close ancestors have been exposed to warmer conditions than the Lithuanian genotypes, particularly during summer, the period coinciding with the onset of new leaf infections (Landolt et al. 2016). Although tree resistance to disease is also expected to vary plastically with temperature (Sturrock et al. 2011), the differentiation in virulence plasticity between the two *H. fraxineus* populations, together with the results of Bécans et al. (2025), is consistent with the hypothesis of thermal adaptation in *H. fraxineus* in southern Europe. Interestingly, such adaptation would have occurred following the severe initial bottleneck experienced by the European *H. fraxineus* population (McMullan et al. 2018), despite potentially strong asymmetric gene flow from core populations into newly founded populations (Gross et al. 2014). Similar patterns of differentiation consistent with thermal adaptation have been reported in plant pathogens in recent years (*e.g*., Robin et al. 2017).

Nonetheless, there are several limitations to the present study that limit the inferences that can be drawn regarding thermal adaptation. First, our reaction norms, based on only two temperatures (22 and 26 °C), may not adequately capture the typically non-linear thermal responses of traits such as *H. fraxineus* virulence (Malusare et al. 2023). This limitation is particularly important because temperature variation across a broader range is likely to affect fungal virulence and ash tree resistance differently (*e.g*. Desaint et al. 2021), which may alter the patterns observed here. Second, with only two populations, our study cannot provide strong evidence for thermal adaptation near the southern edge of the ash dieback expansion front. Sampling additional populations from both northern and southern Europe, or along a latitudinal gradient, will be necessary to determine whether the patterns observed here are consistent across the species’ range and to provide stronger support for the thermal adaptation hypothesis. Third, determining whether the differences in reaction norms between the studied populations result from selection rather than genetic drift will require genomic analyses to quantify neutral divergence and potentially identify trait-associated loci through genome-wide association studies.

## Conclusion

Our study suggests that epidemic severity and disease progression may be greater than anticipated under the warm summer conditions of southern Europe. These findings highlight the importance of accounting for evolutionary changes in pathogen responses to environmental variation when assessing disease risk. Further studies should assess virulence plasticity across a larger number of *H. fraxineus* genotypes and a broader thermal range, while accounting for other environmental factors, such as humidity, that may interact with temperature and host resistance. In parallel, genomic analyses comparing historically and recently established *H. fraxineus* populations, particularly those sampled near the southern front of the ash dieback epidemic, will be essential for further investigating the evolutionary processes underlying variation in virulence plasticity.

## Acknowledgments

Clémence Becans received a doctoral scholarship from the French Ministry for Research and Higher Education. We thank Patrick Léger, Céline Lalanne, Aurélien Kohler, Estelle Ducourtioux and Noémie Chaland for their assistance during the experimental work. We are grateful to Olivier Fabreguettes for his help with the analysis and interpretation of genetic marker peaks. We thank the Phenobois facility (Phenobois, INRAE 2018. Wood and Tree Physicochemical Phenotyping Facility for Genetic Resources, https://doi.org/10.15454/1.5572410490640864E12) for its contribution to NIR spectroscopy measurements. We also thank the Gentyane platform (UMR GDEC, INRAE / Université Clermont Auvergne, Clermont-Ferrand, France) for technical assistance and genotyping services.

## Data availability

The data and script used in this study are available at https://doi.org/10.57745/JG7XSQ (Soularue et al. 2026).

