## Supplementary tables and figures for "Thermal plasticity of virulence differs between two latitudinally distant populations of the ash dieback pathogen"

### Supplementary material

**Table S1.** Genotypic diversity of *Hymenoscyphus fraxineus* isolates from the Italian and Lithuanian populations, based on 18 SSR loci. *N*: number of isolates; *MLG*: number of unique multilocus genotypes; *H*: Shannon-Wiener diversity index; *lambda*: Simpson's index; *E.5*: genotypic evenness; *Hexp*: expected heterozygosity; *Ia* and *rbarD*: indices of association, testing for linkage disequilibrium. Total: both populations combined.

| Population | N | MLG | H | lambda | E.5 | Hexp | Ia | rbarD |
|---|---|---|---|---|---|---|---|---|
| Italian | 15 | 15 | 2.71 | 0.93 | 1 | 0.37 | 0.04 | 0.002 |
| Lithuanian | 15 | 15 | 2.71 | 0.93 | 1 | 0.37 | -0.01 | 0,000 |
| Total | 30 | 30 | 3.40 | 0.9 | 1 | 0.38 | 0.01 | 0,001 |

**Table S2**. Spearman's correlations (*r)* between the observed values of the studied virulence components at each environmental condition througout the experiment.

| Virulence components | Environment | *r* | *p-value* |
|---|---|---|---|
| lesion length *vs* symptom occurrence | 22°C / *F. excelsior* | 0.59-0.74 | **<0.01** |
| lesion length *vs* severe symptom | 22°C / *F. excelsior* | 0.12-0.66 | **<0.04** |
| lesion length *vs* symptom occurrence | 26°C / *F. excelsior* | 0.55-0.74 | **<0.01** |
| lesion length *vs* symptom occurrence | 26°C / *F. excelsior* | 0.49-0.76 | **<0.01** |
| lesion length *vs* symptom occurrence | 22°C / *F. ornus* | 0.69-0.81 | **<0.01** |
| lesion length *vs* symptom occurrence | 22°C / *F. ornus* | 0.35-0.71* | **<0.01** |

* non-significant (*r = 0.04, p = 0.08) at day 42*

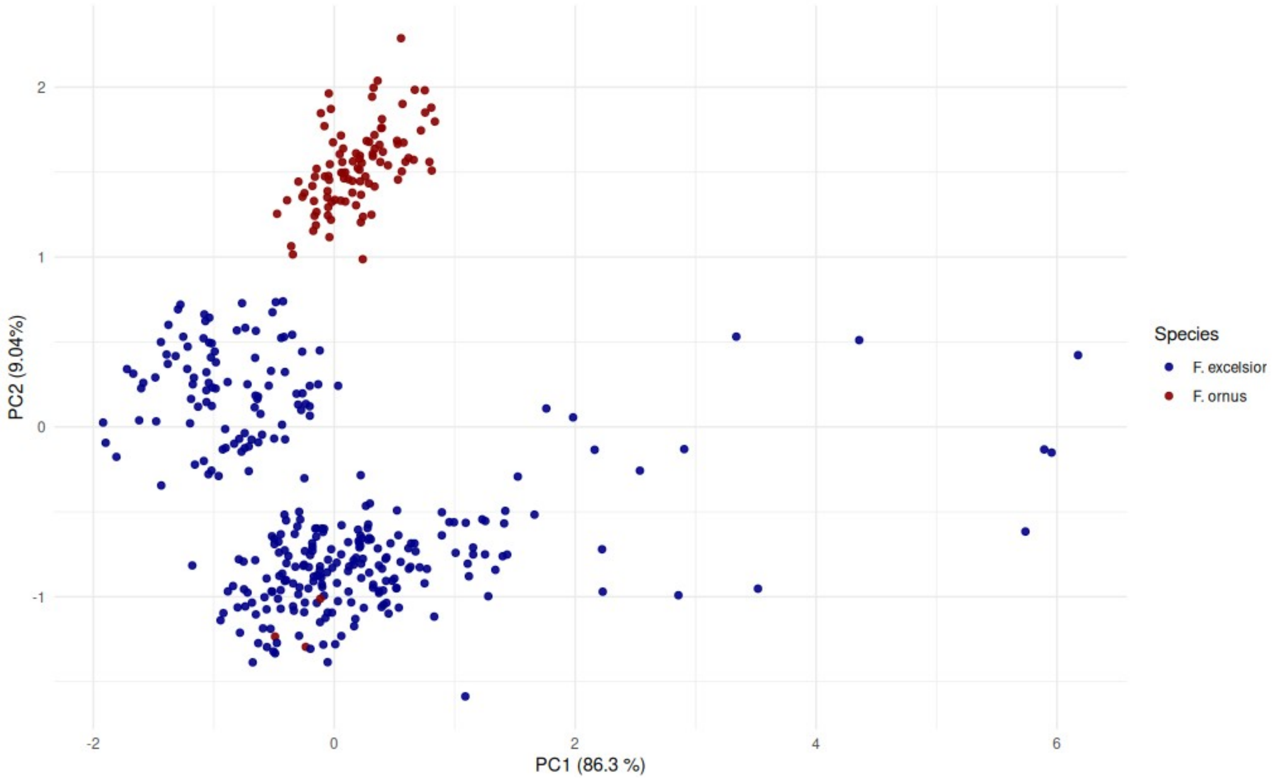


**Figure S1**: Principal component analysis of near-infrared spectral data obtained from dried and ground leaves of *Fraxinus excelsior* (blue) and *Fraxinus ornus* (red) seedlings.

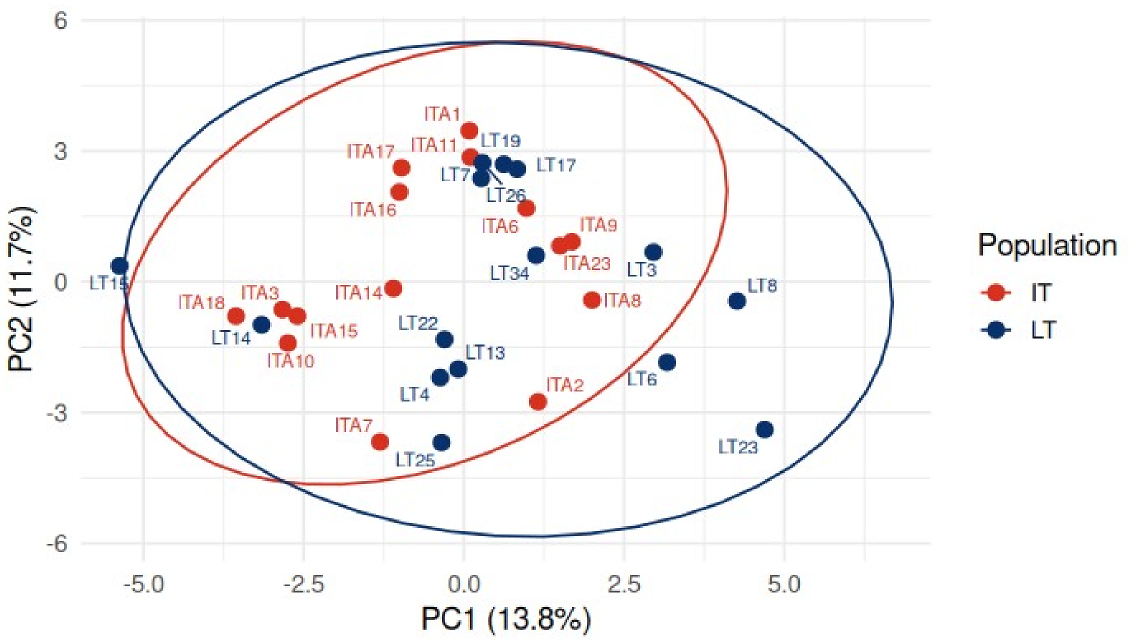


**Figure S2**. Principal Component Analysis (PCA) of *Hymenoscyphus fraxineus* isolates (15 Lithuanian (blue) and 15 Italian (red))) genotyped at 18 SSR loci. Ellipses represent the 95% confidence interval around each population's centroid.

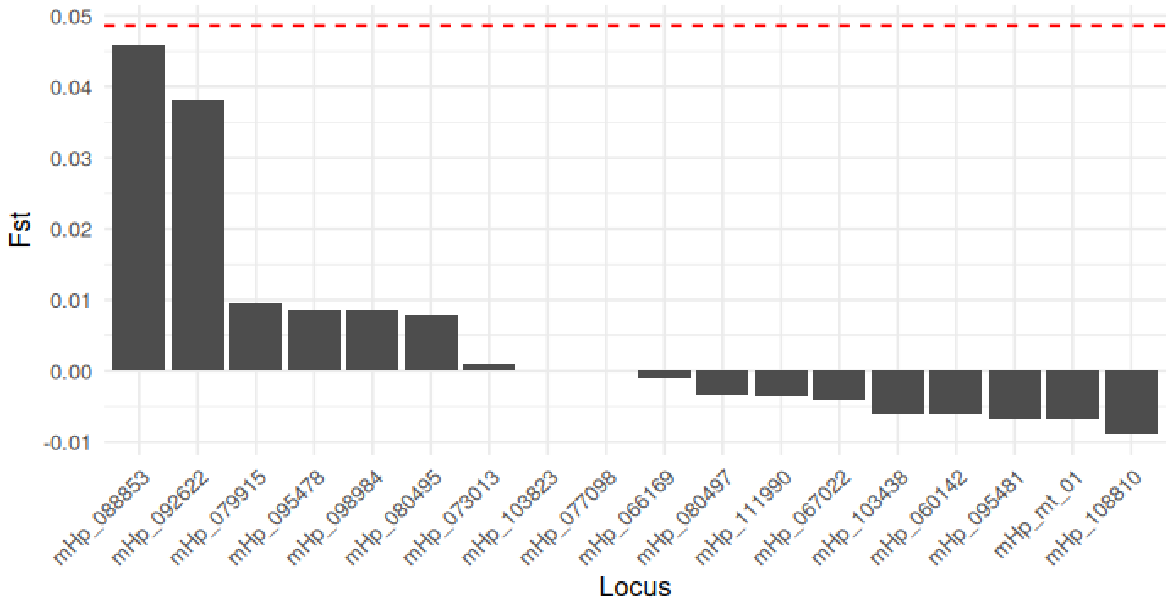


**Figure S3.** Locus-specific Fst estimates (Weir & Cockerham, 1984) calculated for each SSR locus between Lithuanian and Italian *Hymenoscyphus fraxineus* populations. The red dashed line indicates the overall multilocus differentiation estimate.

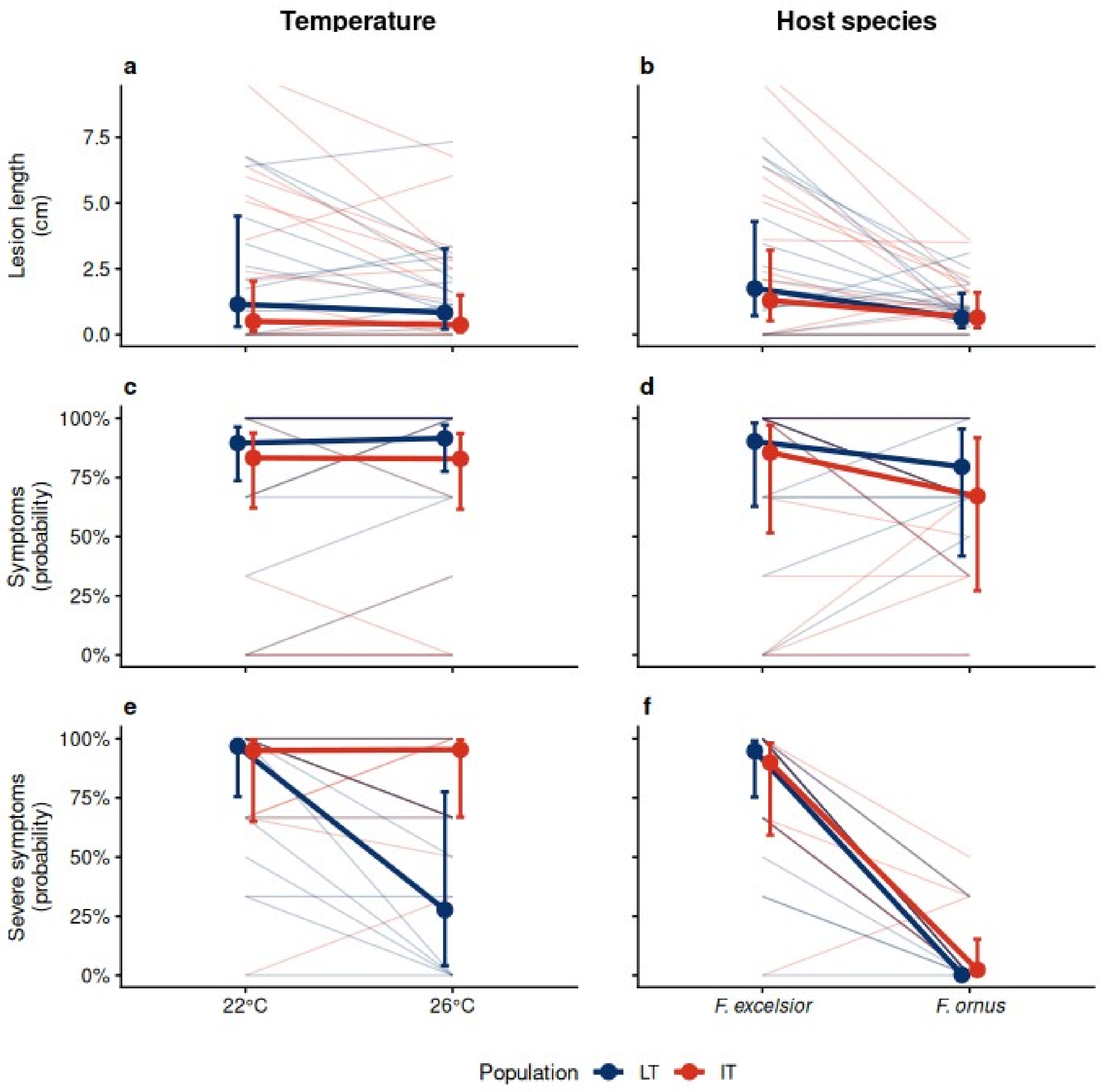


**Figure S4**. Reaction norm estimates for lesion length (cm) (a, b), symptom occurrence (c, d), and symptom severity (e, f) for two *Hymenoscyphus fraxineus* populations (blue: Lithuania, red: Italy, ) in response to temperature (left column) and host species (right column), on day 35 after inoculation. Thin lines represent individual reaction norms, estimated from three replicates per isolate and modality. Thick lines represent the mean reaction norm for each population. Large dots represent the marginal mean estimate value for each population. Error bars indicate 95% confidence intervals of the mean.
